# Intermediate antiparallel fibrils in Aβ40 Dutch mutant aggregation: nanoscale insights from AFM-IR

**DOI:** 10.1101/2023.03.21.533667

**Authors:** Siddhartha Banerjee, Tanmayee Naik, Ayanjeet Ghosh

## Abstract

Cerebral Amyloid Angiopathy (CAA), which involves amyloid deposition in blood vessels leading to fatal cerebral hemorrhage and recurring strokes, is present in the majority Alzheimer’s disease cases. Familial mutations in the amyloid β peptide is correlated to higher risks of CAA, and are mostly comprised of mutations at residues 22 and 23. While the structure of the wild type Aβ peptide has been investigated in great detail, less is known about the structure of mutants involved in CAA and evolutions thereof. This is particularly true for mutations at residue 22, for which detailed molecular structures, as typically determined from Nuclear Magnetic Resonance (NMR) spectroscopy or electron microscopy, do not exist. In this report, we have used nanoscale infrared (IR) spectroscopy augmented with Atomic Force Microscopy (AFM-IR) to investigate structural evolution of the Aβ Dutch mutant (E22Q) at the single aggregate level. We show that that in the oligomeric stage, the structural ensemble is distinctly bimodal, with the two subtypes differing with respect to population of parallel β-sheets. Fibrils on the other hand are structurally homogeneous, with early-stage fibrils distinctly anti parallel in character, which develop parallel β-sheets upon maturation. Furthermore, the antiparallel structure is found to be a persistent feature across different stages of aggregation.

## Introduction

Aggregation of amyloid β proteins into oligomeric species and subsequently fibrils is known to be the major molecular pathway driving the development of Alzheimer’s disease (AD)^1-3^. Sequential proteolytic cleavage of amyloid precursor protein (APP) by β and gamma secretase results in a series of peptide fragments typically with 39-43 residues, which aggregate into plaques in the cerebral cortex of AD patients. Another pathological marker of AD is Cerebral Amyloid Angiopathy (CAA), which is characterized by fatal cerebral bleeding and recurrent strokes, caused by amyloid deposition primarily in the cerebral blood vessels^4-6^. Two types of AD have been identified based on the age of disease onset and genetic predisposition^7-8^. Most common form of AD is sporadic which occurs in individuals after the age of 65 years and characterized by slow progression of the disease. CAA is present in over 90% cases of non-familial age-related Alzheimer’s disease. The other type is familial which is rare and found at an earlier age with higher severity. Most of familial AD is associated with mutation at residue 22, including Arctic (E22G), Italian (E22K) and Dutch (E22Q). Another familial mutation occurs at residue 23, Iowa (D23N). Patients with familial mutations are at higher risk of CAA^7, 9^. Hereditary Cerebral Hemorrhage with Amyloidosis-Dutch type (HCHWA-D), which causes severe CAA along with strokes and early onset dementia^10^, originates from a point mutation at codon 693 of APP producing a single mutation of Glu to Gln at residue 22 resulting in the Dutch mutant (E22Q) of Aβ. However, compared to wild type Aβ^11-13^, significantly less is known about the familial mutants. *In vitro* experiments show that E22Q mutant aggregates slightly faster compared to wild type Aβ, and mature fibrils of the Dutch and Iowa mutants have been shown to have a predominantly parallel cross β structure^14-18^, which are believed to be generated from antiparallel prefbrillar aggregates. However, while detailed structural models of the Iowa mutant have been generated through Nuclear Magnetic Resonance (NMR) spectroscopy, such structures for the D22 mutants are yet to be determined. Furthermore, in the context of the role of Aβ in CAA and AD, it is now believed that early-stage aggregates play a more significant role than mature fibrils^19-20^, which are significantly harder to isolate and structurally characterize. Limited structural information is hence available about these early-stage aggregates that mediate the formation of mature fibrils. Specifically, how the structure of Aβ D22 mutants evolves during several stages of aggregation is yet to be fully understood. This information is critical since amyloid proteins in AD brain are expected to exist in several aggregated forms namely oligomers, protofibrils and fibrils at any given time, and understanding the structural ensemble can potentially unlock therapeutic possibilities that target vascular amyloid aggregates, which currently do not exist.

In this report, we have applied atomic force microscopy (AFM) augmented with infrared spectroscopy (IR) to investigate the structural evolution of E22Q mutant throughout the aggregation process, wherein AFM generates high-resolution topographic image of the aggregates, while IR provides structural information^21-22^. AFM-IR thus simultaneously provides both morphological and structural insights, allowing for spectra and hence structures to be mapped to individual aggregates and resolving heterogeneities in the ensemble^23-24^, which is difficult to obtain from conventional spatially averaged techniques such as NMR and Fourier Transform Infrared (FTIR) spectroscopies. We demonstrate that E22Q mutant forms oligomers with two distinct structural distributions, both containing antiparallel structure. Fibrils on the other hand are more homogeneous; however, early-stage fibrils have distinct anti-parallel character which eventually develop parallel β structure. All fibrils have persistent anti-parallel β-sheets, which is not transient, contrary to previously known structures of Aβ mutants like Iowa, where anti-parallel fibrils were metastable. These results underscore the necessity for investigating the structure of amyloid aggregates from a point of view where the evolution of the structures is looked at as a dynamic process rather than a static system.

## Materials and Methods

### Preparation of Dutch Aβ40 mutant aggregates

Lyophilized Dutch Aβ40 (Anaspec, USA) was initially treated with 1,1,1,3,3,3-hexafluoroisopropanol (HFIP) to remove any preformed aggregates. HFIP treated solution was kept at room temperature for 15 minutes and then evaporated by keeping the vial under vacuum desiccator. Aggregation of mutant was carried out with a protein concentration of 100 µM in 10 mM phosphate buffer (Sigma-Aldrich, USA), pH 7.4 at 37ºC without agitation.

### Sample preparation for AFM-IR experiment

Aliquots were taken out from the aggregation mixture at 2h, 8h, 36h, 72h and 192h (8 days). Then the solution was half-diluted and a 5 µL droplet was deposited on to ultraflat gold substrates (Platypus Technologies, USA). The droplet was incubated for 5 minutes at room temperature in a covered petri dish and then rinsed with 50 µL of Milli-Q water. Then the substrate was dried with gentle stream of nitrogen gas.

### AFM-IR experiment

AFM-IR experiments were performed with Bruker NanoIR3 (Bruker Corporations, USA) equipped with a mid-IR quantum cascade laser (MIRcat, Daylight solutions, USA) at rom temperature with low relative humidity. AFM topographs and IR spectra were collected in tapping mode. The resonance frequency and the spring constant of cantilevers were 75 ± 15 kHz and 1–7 N/m, respectively. Scan rate for the AFM imaging was kept at 1.0 Hz. The experiment was carried out by first acquiring a high-resolution AFM topograph and then the tip was positioned on oligomers/fibrils to obtain the IR spectrum. IR spectral resolution was 2 cm^-1^ and total 128 co-additions at every point with 64 co-averages for each spectrum were applied. Multiple AFM topographs were acquired in between spectral scans to mitigate effects from thermal drifts. The total number of spectra acquired for different time points are as follows: 30 (2 h), 35 (8 h), 24 (36 h, 72 h and 8 days each).

### Data analysis

AFM images were minimally processed with Gwyddion software. IR data were processed using the MATLAB software package by applying (3, 7) Savitzky–Golay filter and a 3-point moving average filter. A baseline correction was applied for each spectrum. Mean spectra were fitted with four Gaussians peaks and the details are mentioned in Supporting Information.

## Results and discussions

To understand the structure of Aβ40 dutch mutant (E22Q) aggregates and their evolution at different stages of aggregation, the E22Q mutant peptide was incubated in 10 mM phosphate buffer (pH 7.4) at 37°C under quiescent conditions. Oligomers were observed within 2 h of incubation in the AFM topographs as shown in Fig. 1A. The presence of a film of small oligomeric aggregates along with isolated larger oligomeric species can be seen in the AFM image. The IR spectra in the amide I region, recorded from these oligomeric species, can be classified into two main subtypes; the mean spectrum corresponding to each subtype is shown in Fig. 1B and C.

**Figure 1.**
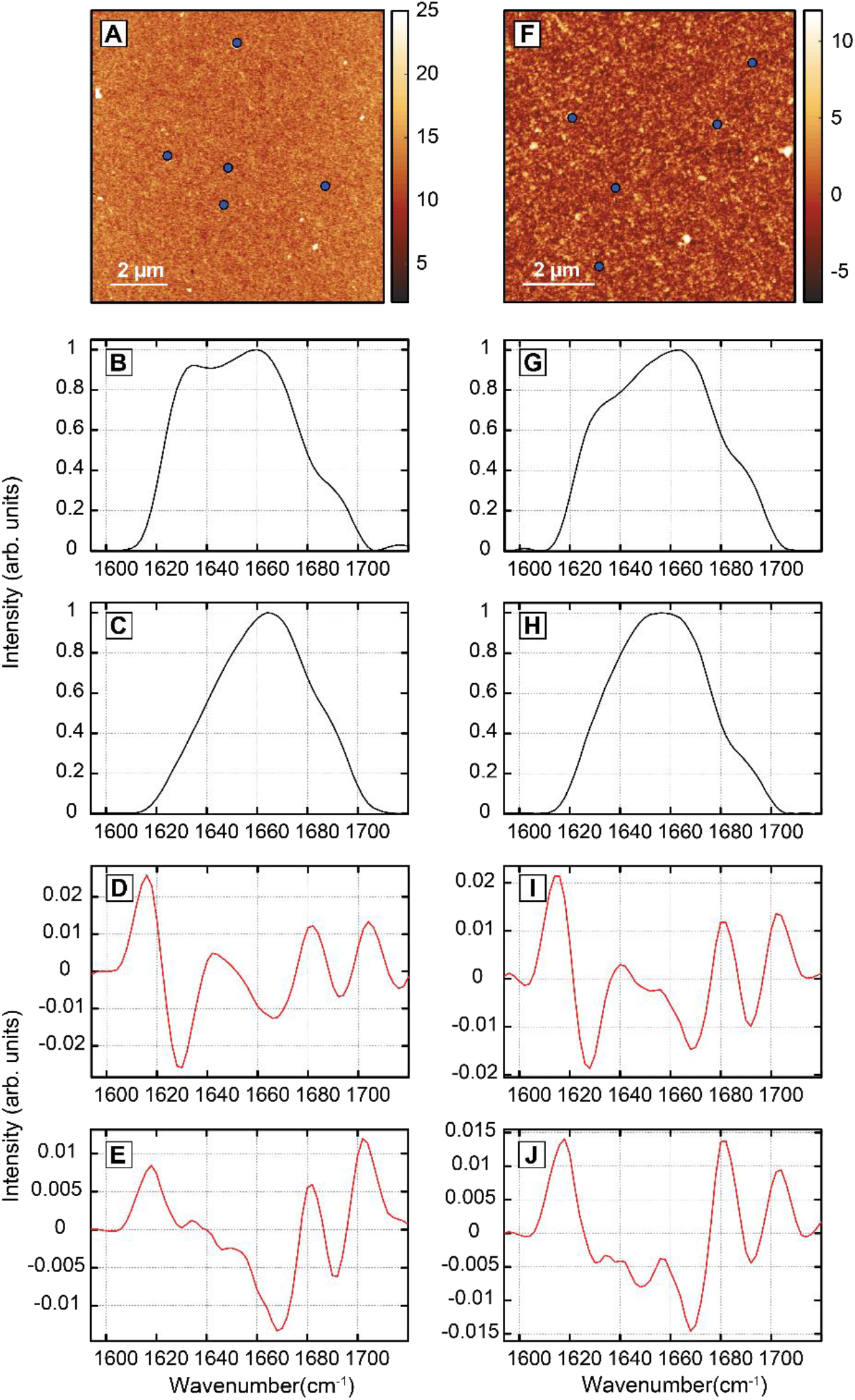
AFM topographs and IR spectra of Aβ Dutch mutant oligomers. (A) AFM image of oligomers after 2 h. (B-C) Mean spectra of the two spectral types for 2 h oligomers. (D-E) Second derivatives of the mean spectra of 2h oligomers. (F) AFM image of 8 h oligomers. (G-H) Mean spectra of the two oligomeric subtypes. (I-J) Second derivatives of mean spectra for 8 h oligomers. Representative spatial location from which spectra were acquired are indicated with blue circles.

Representative spectra are shown in Figure S1. The first type shows two distinct peaks at 1634 cm^-1^ and 1666 cm^-1^ with a shoulder at 1690 cm^-1^ (Fig. 1B). The other type exhibits the same set of constituent bands, and differs from the first with respect to the relative intensities of these bands: the peak at 1666 cm^-1^ is the most intense in this case, while the 1630 cm^-1^ band is much less prominent (Fig. 1C). The differences can be better discerned from the spectral second derivatives, shown in Fig. 1D-E. The shoulder at 1690 cm^-1^ is persistent in both the spectral subtypes. The first type is less abundant in comparison to the second and constituted ∼27% (8/30) of all spectra acquired. It is important to note in this context that the spectra are not meant to indicate a specific statistical distribution, and the spectral abundances of the subtypes should not be interpreted as exact. It is well known that amide I vibrations are reflective of protein secondary structure, and thus presence of two distinct spectral types suggests structural variation within the oligomeric aggregates. The spectra thus demonstrate that early-stage oligomers of the Dutch mutant are structurally heterogeneous and do not conform to a singular secondary structure. This is consistent with previous reports on wild-type Aβ^24^. An amide I peak centered at ∼1630 cm^-1^ indicates the presence of ordered parallel/antiparallel β-sheet arrangement and an additional peak at ∼1690 cm^-1^ can be attributed to the anti-parallel β-structure, whereas a peak at ∼1666 cm^-1^ can originate from both random coil and β turns^25-26^. The absorption at all these three regions with varying intensities suggest that Dutch mutant oligomers formed at early stages of aggregation adopt conformations with a combination of parallel, anti-parallel β structure and random coil/β turn structural elements. The second derivative spectra of one of the oligomeric types indicate the presence of an additional band at ∼1648cm^-1^, which can arise from disordered random coils or from helical structures^25-26^. We discuss the structural implications of the spectra later in the manuscript.

After 8 hours of aggregation, oligomers were still observed (Fig. 1F). IR spectra were recorded from different oligomers as before, which displayed similar trends as the 2 h aggregates. The spectra can again be grouped into two classes that differ with respect to the relative intensity of constituent bands. The average spectra for each class and the corresponding second derivatives are shown in Fig. 1G-J. The peak of the amide I band remains at 1666 cm^-1^, whereas the intensity of two shoulders at 1634 cm^-1^ and 1690 cm^-1^ changes among different oligomers. This suggests that the structural heterogeneity observed in earlier oligomers is not a transient feature and the structural ensemble does not narrow significantly in the oligomeric stages of aggregation, i.e., oligomers at different stages of aggregation exhibit similar heterogeneity. The peak/band at ∼1648cm-1 is also present in the second derivative spectra for both types, similar to the 2 h aggregates, further indicating little change in the oligomeric structure at this stage of aggregation. Interestingly, we do observe a change in the relative abundances of the oligomer types: the one exhibiting higher intensity at 1630cm^-1^ is more abundant at 8 h, constituting nearly 65% of spectra (23/35).

Fibrillar aggregates started appearing after 36 h of incubation. Fibrils with different lengths are observed on the AFM topographs (Fig. 2A). IR spectra were acquired from different spatial locations along the length of multiple fibrils. The mean fibril spectrum thus obtained from four different fibrillar aggregates is shown in Figure 2B along with the spectral derivatives in Fig. 2C. Additional spectra of fibrils are shown in Figure S2. The fibril spectra exhibit similar spectral features as the oligomers: the peak of the amide I band remains at 1666 cm^-1^ with two shoulders at 1630 cm^-1^ and 1692 cm^-1^ (Fig. 2B) and a small peak at 1650cm^-1^ that is only apparent in the second derivatives (Fig. 2C). However, the shoulder at 1634cm^-1^ is not as prominent as some of the oligomeric spectra, and in fact the overall spectra are similar to one of the oligomeric subtypes. The fibril spectra also do not exhibit the distinct differences observed for oligomers and point towards a more homogeneous structural distribution. After 72 h, more fibrils were observed in the AFM topographs along with fibrillar networks (Fig. 2D). The mean spectrum from four different fibrillar aggregates, shown in Fig. 2E, and the corresponding second derivative (Fig. 2F) are very similar to the fibrils which are formed after 36 h, and exhibit small differences in intensities at 1630 cm^-1^ and 1692 cm^-1^. The aggregation was followed up to eight days (192 h) to identify any significant structural modifications of the aggregates due to maturation. AFM topographs show the presence of fibrillar networks and some isolated clusters of fibrils after eight days (Fig. 2G). Morphologically, individual fibrils seen after 8 days of maturation do not show any major deviation from those observed earlier, except they become clustered as appeared in AFM topographs (Fig. 2G). However, fibrillar spectra do exhibit significant differences. The average spectrum and its second derivative from these fibrillar species is shown in Fig. 2H-I. The spectra appear to be composed of the same constituent bands, with most intense component centered at 1666 cm^-1^, and presence of two shoulders at 1630 cm^-1^ and 1692 cm^-1^. The major difference observed in these spectra is that the intensity of the 1630 cm^-1^ shoulder is significantly higher compared to the previous time points (Fig. 2A-F), indicating the increase in parallel β-sheet structure. Interestingly, for these fibrillar aggregates, we do not observe distinct spectral subtypes either, and the increase in the relative population of the 1630 cm^-1^ band happens in all fibrillar aggregates.

**Figure 2.**
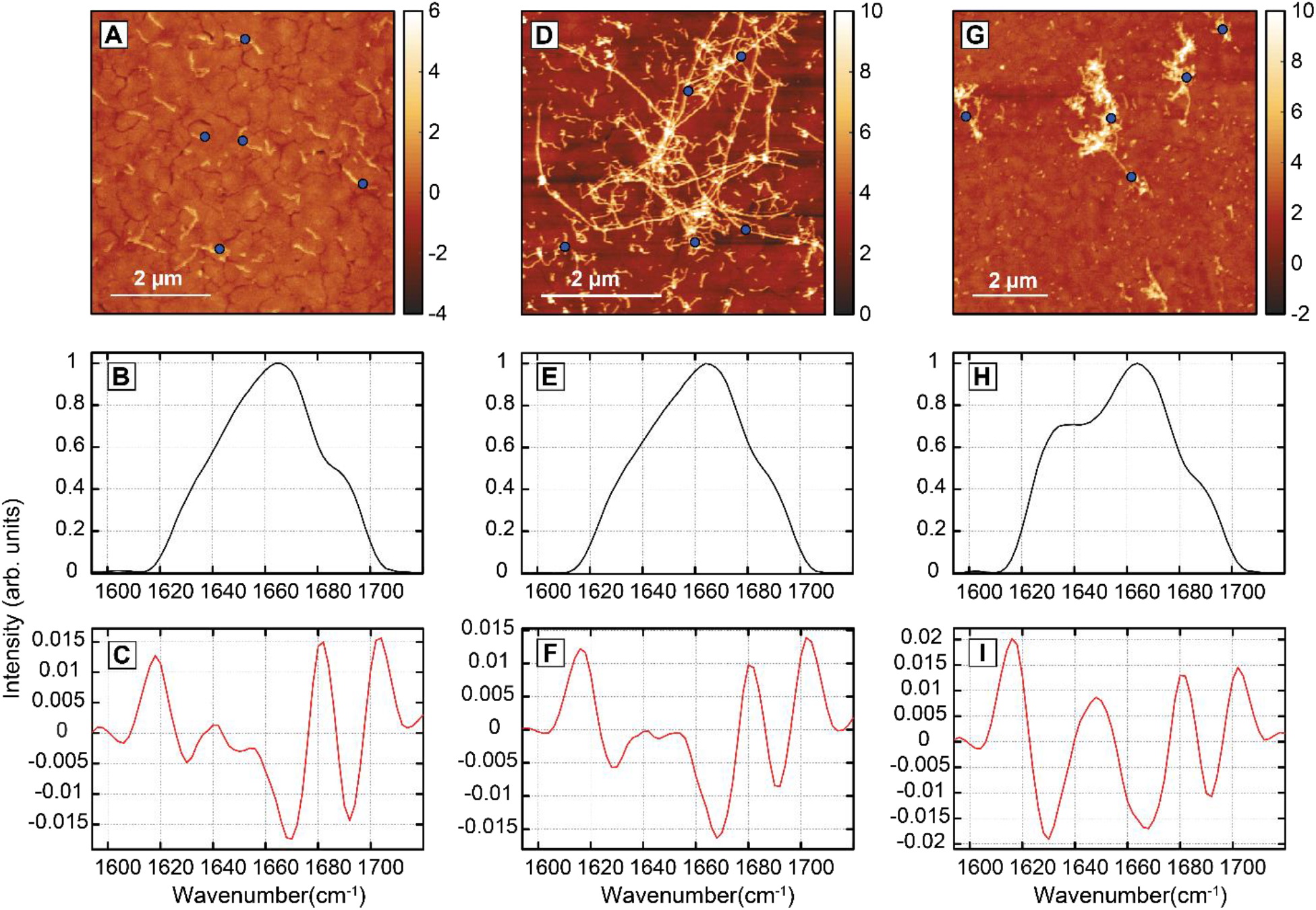
AFM topographs and IR spectra of Aβ Dutch mutant fibrils. (A) AFM image of fibrils after 36 h. (B-C) Mean spectrum and spectral second derivative for 36 h fibrils. (D) AFM image of 72 h fibrils. (E-F) Mean spectra and corresponding second derivative of 72 h fibrils. (G) AFM image of 8-day fibrils, and corresponding mean spectrum and second derivative (H-I). Representative spatial locations of spectra are indicated with blue circles.

These results shine light on the structural evolution of the Aβ40 Dutch mutant (E22Q) at the single aggregate level during aggregation, starting from early-stage oligomers to mature fibrils. The presence of two distinct spectral types at the early stages of aggregation (2 h and 8 h) indicate that the oligomers do not conform a singular structure. These structural variations do not appear to be correlated with any morphological difference between oligomers. This heterogeneity underscores the need for spatially resolved measurements for investigating structure of amyloid aggregates: the average spectra typically acquired in conventional approaches like FTIR cannot be easily attributed to the presence of multiple conformations of specific aggregate species in the ensemble. Both the oligomeric subtypes exhibit the same set of sub-bands at 1630 cm^-1^, 1666 cm^-1^ and 1692 cm^-1^ respectively, as can be seen from the spectra and their second derivatives (Fig.1D-E and I-J). The amide I infrared spectra of peptides and proteins is well known, and a peak at 1620-1630 cm^-1^ is typically attributed to β-sheet secondary structure, whereas a band at 1666 cm^- 1^ can arise from random coils and/or β turns^25-26^. However, the 1630 cm^-1^ band is present in both parallel and antiparallel β-sheets, with the latter exhibiting an additional peak at ∼1690 cm^-125-26^. Given that the spectra observed herein exhibit both aforementioned bands, we can conclude that both types of oligomers contain antiparallel β structure. In fact, the key difference between the oligomeric spectral types is the relative intensity of the 1630 cm^-1^ peak, whereas the intensity of the 1690 cm^-1^ peak remains virtually unchanged. Since the latter arises only from antiparallel β structure, while the former can be attributed to both and antiparallel, we conclude that the oligomeric subtypes differ with respect to their parallel β structure. However, this does not preclude the presence of parallel β-sheets from certain oligomers or imply that one subtype only contains antiparallel β-sheets only while the other contains both parallel and antiparallel; the spectra are instead reflective of the relative abundance of the two types of β structure. Essentially, the spectra indicate that one subtype contains more parallel structure than the other. For clarity, we will refer to these subtypes as type 1 and type 2 oligomers respectively for the rest of this manuscript. The spectra type 2 oligomers for 2 h and of both the subtypes for 8 h oligomers additionally have a small sub-band at ∼1648 cm^-1^. A peak at ∼1645-1650 cm^-1^ for proteins is typically attributed to alpha helical structure^26^; however, it can also arise from disordered random coils. In fact, the reported spectral ranges for random coil absorption overlaps with helices but also to some degree with β turns^26^, which makes unequivocal attribution of the band at 1648 cm^- 1^ difficult. Given that we observe a band at 1666 cm^-1^ that is typically attributed to random coils in infrared spectra of amyloid aggregates^18, 27-28^, the band at 1648 cm^-1^ likely arises from a different structural element. Silva et. al. have shown that early-stage fibrils of short peptides derived from the prion protein involve structural disorder in the form of strand misalignments, leading to out-of-register β-sheet configurations, which can have a blueshifted infrared signature^27^. It is possible that that the apparent helical band observed here arises from similar register defects in the aggregates. However, most reported infrared spectra of amyloid aggregates correspond to solution phase measurements, and it is well known that changing or removal of the electric field arising from the solvent molecules can alter the peak positions significantly^29-30^. This makes direct comparisons between previously reported spectral parameters and the features observed here challenging, and elucidating the precise structural origin of the 1648 cm^-1^ band thus requires additional experimental metrics, such as those afforded by isotope edited measurements. We aim to address this in future work. It should also be noted in this context that AFM-IR spectra differ fundamentally with bulk isotropic FTIR measurements and can depend on the laser polarization and illumination configurations used^22, 31^. For amide I spectra, while the overall spectrum is still constituted by the same underlying sub-bands, the contribution of certain bands can be amplified over others. Thus, the relative intensity of specific components, as typically observed from bulk FTIR measurements, may not be exactly reflected in AFM-IR spectra. However, since all the data are acquired under identical conditions, the variations in intensities of specific peaks between different spectra nevertheless reflect a change in the structural distribution, which in our case, points to a key conclusion: the oligomeric aggregates of the Dutch mutant, while structurally divergent, always contain antiparallel β-sheet structure.

For fibrils that appear after 36-72 h of aggregation, we observe only one spectral type. This is verified by acquiring spectra from multiple fibrillar aggregates (Fig. S2). The fibril spectra closely resemble those of the type 2 oligomers, i.e., those with more relative antiparallel conformation. This indicates that: a. early-stage fibrils have distinct antiparallel character, and b. they also have significant amount of inherent structural disorder (i.e., presence of ordered β-sheets and disordered secondary structural elements from the same spatial location), as evidenced by the peaks at 1648 cm^-1^ and 1666 cm^-127-28^. For fibrils observed after 8 days of aggregation, we find the 1630 cm^-1^ peak to be more intense compared to 36-72 h aggregates, indicating that with maturation, the fibrils develop more ordered β-sheets, which is expected and consistent with the general picture of amyloid aggregation. These β-sheets are likely parallel in character since the intensity of the 1692 cm^-1^ peak does not exhibit any major change. This also implies that fibrils, even after maturation, still contain some amount of antiparallel structure. In fact, we find that at all stages of aggregation, the Dutch mutant structure involves antiparallel β-sheets. This is a marked difference compared to wild type Aβ^11-12, 32-33^.

Taken together, we observe two distinct types of oligomers between 2-8 hours: with mostly antiparallel character (type 2) and with parallel and antiparallel character (type 1), fibrils that spectrally resemble only one subtype of oligomers (type 2) between 36-72 hours, and finally fibrils after 8 days that are spectrally similar to the other type of oligomers (type 1). The similarity in spectra between the early (36-72 h) fibrils and antiparallel oligomers (type 2) suggests that the former evolves from the latter. Since we do not observe any fibrils between 36-72 hours that spectrally resemble type 1 oligomers, it leads to the conclusion that the antiparallel oligomers possibly evolve into fibrils faster than their parallel (type 1) counterparts. The parallel oligomers are kinetically hindered in comparison: they need to undergo structural reorganization to form fibrils, as evident from the spectral difference between the two species. Alternatively, the parallel oligomers can be completely off pathway, and never undergo the structural reorganization, thus never converting to fibrils at all. The relative abundance of each of the oligomeric types between 2 and 8 hours offers additional insights in this context. While the abundances do not necessarily reflect the exact population of the oligomeric subtypes, we do find a notable increase in the relative abundance of the type 1 oligomers after 8 hours. This can be explained if the type 2 oligomers, which were more abundant at 2 h, have started converting into fibrillar aggregates, leading to the overall ensemble being skewed towards type 1 aggregates. Alternatively, a greater abundance of type 1 oligomers could mean that type 2 oligomers are structurally converting into type 1, which then would need to undergo reorganization themselves to form the fibrils we see at 36-72 h. We reject this possibility as it is less likely that oligomers would undergo two steps of structural reorganization to form fibrils that are similar to their initial structure. After 8 days we do not observe any fibrils that spectrally resemble the type 2 oligomers or the 36-72 h fibrils. One explanation of this is that these fibrils are formed through maturation of earlier fibrillar aggregates, which all convert to the fibrillar structure seen after 8 days. In that event, we can expect to see an overall increase in the population of β-sheets compared to 36-72 fibrils, which is known to be a structural hallmark of fibril maturation. The other possibility that needs to be considered is that the 8-day fibrils evolve from the type 1 oligomers. This would imply that the 36-72 h fibrils are off pathway. In that event we can expect to observe two types of fibrils in the ensemble, those that have formed from the type 1 oligomers, and those from 36-72 hours that haven’t matured any further. However, we only see one subtype of fibrils, which makes the above possibility less likely. Additionally, if the 8-day fibrils are indeed evolved from the parallel oligomers, their conformational distribution should at least reflect more order than the latter, i.e., they should have higher/similar relative β-sheet character than the parallel oligomers, but not less. The spectral interpretation above relies largely on difference in spectral intensities at specific wavenumbers combined with second derivative analysis of spectra. While the latter can be indicative of presence of underlying peaks in spectra, it does not allow for quantitative assessment of secondary structural distributions. To get further insights into the relative populations of the structural components at each stage of aggregation, we therefore employed spectral fitting to deconvolute the fibril and oligomeric mean spectra. The number of peaks and their positions, as observed in the second derivative spectra, were used as a starting point for fitting. It was observed that all spectra are best fit to four peaks. The fitted spectra are shown in Figure 3A-G. The fitting procedure is detailed in the Supporting Information, and the fit parameters are listed in Table S1. From the spectral fits, we calculated the β-sheet population, which we define as the area under the curve (AUC) of the β-sheet peak at 1630 cm^-1^. Since this peak can arise from both parallel and antiparallel β-sheets, the AUC represents the overall β percentage of β-sheets in the spectra. On the other hand, the AUC of the 1692 cm^-1^ peak represents the antiparallel β-sheet population only. The results, shown in Figure 3H, reveal interesting insights into the spectral evolution. We see that the relative population of overall β-sheets increases from the 36-72 h to the 8-day fibrils while the antiparallel β structure decreases, consistent with structural ordering upon maturation and with formation of parallel structure in mature fibrils^12^. However, we do not observe a distinct increase in β-sheet character between the parallel oligomers (type 1 of 2 h and 8 h) and 8-day fibrils, and the oligomers actually contain a higher percentage of β-sheets. This strongly suggests that the 8-day fibrils result from structural maturation/evolution of the 36-72 fibrils which in turn evolve from the antiparallel oligomers, and not from the parallel oligomers, as we expect an increase in β-sheet character with time. This means that the parallel oligomers do not readily evolve into fibrils and need to undergo structural reordering prior to fibril formation, and suggests that not all oligomeric species with high β-sheet population are suited to transition into the fibrillar stage.

**Figure 3.**
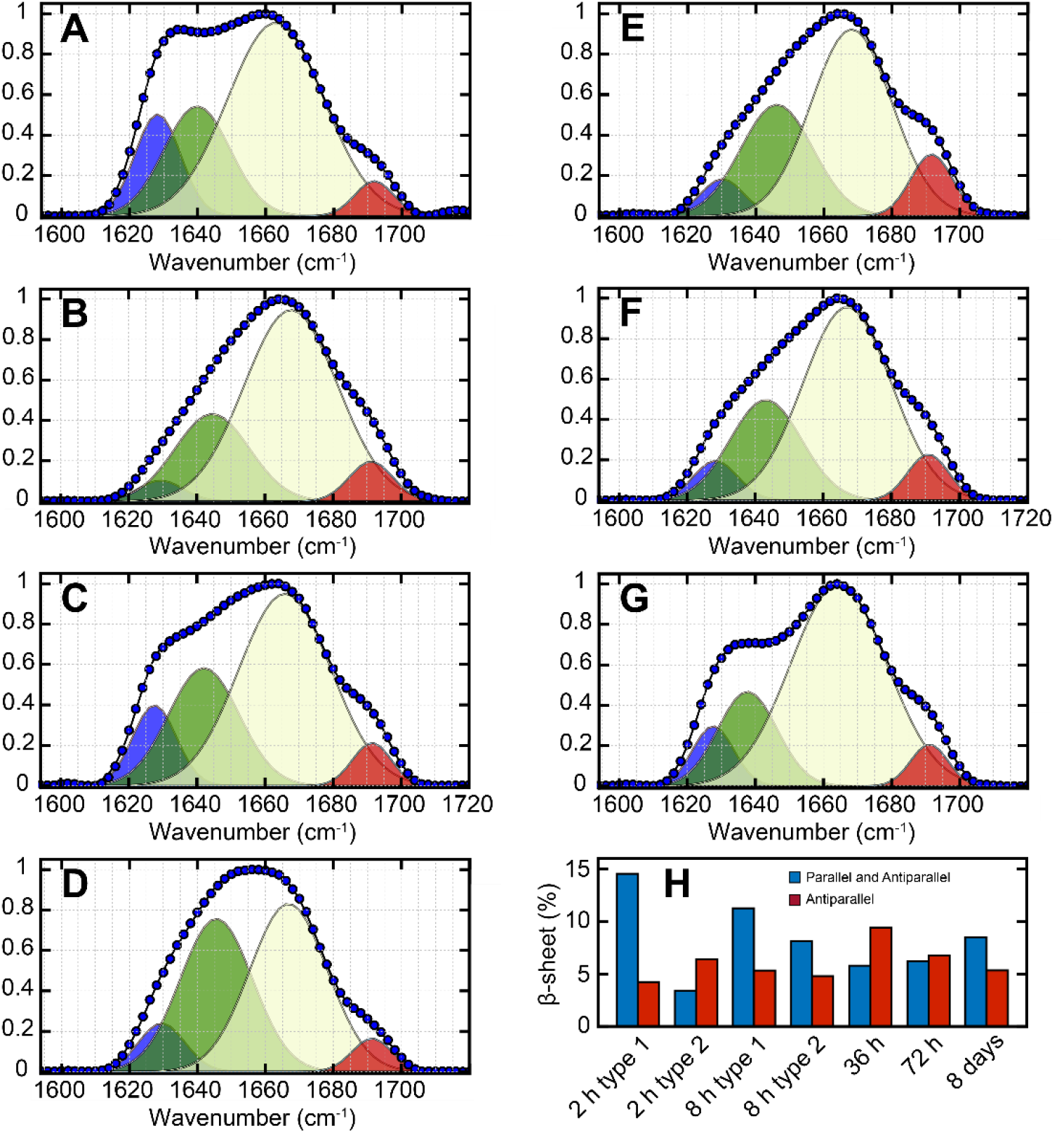
Spectral fitting of Dutch mutant aggregates. Spectral fits of mean spectra of aggregates at (A-B) 2 h, (C-D) 8 h, (E) 36 h, (F) 72 h and (G) 8days. (H) Distribution of β-sheet component in oligomers and fibrils.

Hence the maturation of oligomeric species into fibrils for the Aβ Dutch mutant proceeds through intermediate fibrillar aggregates that have more relative antiparallel character than mature fibrils: essentially the structural transition from antiparallel to parallel occurs in the fibril phase and not in oligomers. We have previously demonstrated that similar structural transitions can occur in smaller wild-type Aβ fragments^34^; the findings reported here point to secondary structural reorganization in fibrils as a more ubiquitous trait of amyloid aggregation. The overall and antiparallel β-sheet populations also increase between the 2 h type 2 oligomers and the 36-72 h fibrils, which is expected if the former were to evolve into the latter. Interestingly, the type 2 oligomers at 8h contain more overall β-sheets than the subsequent 36-72 h fibrils, suggesting that they may also undergo structural reorganization prior to fibril formation that results in loss of β structure. While fibril formation through initial structural reordering of oligomers has been known in the context of amyloid aggregation^27, 33, 35-37^, our findings unfortunately cannot unequivocally validate this mechanism, which requires additional measurements at different aggregation time points so that oligomers at different stages of this structural transition can be arrested and identified in AFM-IR. We hope to address this in future work. Finally, the relative population of antiparallel β-sheets shows a slight increase from oligomers to fibrils, indicating that the antiparallel structure is not transient and present at all stages of aggregation, in agreement with conclusions drawn from spectral intensities. Previous reports^14, 18^ have indicated a parallel β-sheet structure for mature Dutch mutant fibrils; but the structures of early stage and/or intermediate fibrillar species is not known. The Iowa mutant of Aβ has been shown to form metastable antiparallel fibrils under certain aggregation conditions^16-17^, which eventually convert to parallel upon maturation. Our findings mirror these observations and suggest that there are fundamental similarities to the aggregation pathway of Aβ familial mutants. On the other hand, while NMR predicts a parallel cross β structure for mature Iowa mutant fibrils^16^, we see signatures of both parallel and antiparallel β-sheets after 8 days in the Dutch mutant. While this can be a point of divergence in the aggregation mechanism between the two mutants, it is also possible that upon seeded growth over multiple generations, the parallel structure dominates, and the antiparallel character becomes effectively negligible.

## Conclusions

In summary, our results reveal key insights into the structural evolution of the Aβ Dutch mutant peptide from oligomers to fibrillar species at the single aggregate level. The importance of this work lies in the fact that compared to wild-type Aβ, significantly less is known about the structure of its familial mutants. The detailed NMR structure of the Dutch mutant fibrils is yet to be determined, which is the *de facto* gold standard in amyloid structural biology. Of course, even when fibril structures are known, the structural evolution cannot be retraced back to oligomeric and/or prefibrillar species. Recent studies have used FTIR and NMR spectroscopies to determine that early-stage aggregates of the Dutch mutant contain significant antiparallel structure, which is, however, transient in nature and converts to parallel β-sheets in mature fibrils. Our results are in broad agreement with these findings: we do observe a conversion of largely antiparallel oligomers to fibrils with higher parallel character. However, the key insight obtained from this work is that this transition is mediated through an intermediate fibril structure: essentially the fibrils undergo structural evolution and develop more parallel β-sheets with maturation. Furthermore, we also find that the initial ensemble is heterogeneous, with structurally different oligomers present at early stages of aggregation, of which only those with more relative antiparallel character transition to fibrils. Finally, we note that the antiparallel character in Dutch mutant aggregates, under our experimental conditions, is persistent, and shows little variation between oligomers and fibrils; it is the parallel β structure that exhibits significant change. This has not been observed/shown for any of the familial mutants involving a mutation of the D22 residue to the best of our knowledge. For the Iowa mutant of Aβ, solid state NMR studies have shown that early-stage fibrils can be highly polymorphic and can have both antiparallel and parallel β-sheets. Our results are consistent with these reports and point to an overall similarity in the structural distribution of aggregates of the Aβ familial mutants. Taken together, our findings provide novel insights into the evolution of the structural ensemble of the Dutch mutant. Amyloid aggregates in CAA are expected to exist in a range of aggregation states from oligomers to mature fibrils, and hence to develop any drugs that aim at targeting these aggregates, we need to understand the structures of all the members of the dynamic ensemble. The results reported herein address precisely this gap in knowledge and can potentially aid in the development of therapeutic measures that mitigate the symptoms and risks of CAA.

## Supporting Information

The Supporting Information is available free of charge at Representative IR spectra of oligomers and fibrils, details of fitting parameters for IR spectra (PDF)

## Acknowledgement

This research was supported by the National Institutes of Health, grant number R35 GM138162 to A.G.

## References

1. Murphy, M. P.; LeVine, H., 3rd. Alzheimer’s disease and the amyloid-beta peptide. J Alzheimers Dis 2010, 19 (1), 311–323.

2. DeTure, M. A.; Dickson, D. W. The neuropathological diagnosis of Alzheimer’s disease. Molecular Neurodegeneration 2019, 14 (1), 32.

3. Perl, D. P. Neuropathology of Alzheimer’s Disease. Mount Sinai Journal of Medicine: A Journal of Translational and Personalized Medicine 2010, 77 (1), 32–42.

4. Greenberg, S. M.; Bacskai, B. J.; Hernandez-Guillamon, M.; Pruzin, J.; Sperling, R.; van Veluw, S. J. Cerebral amyloid angiopathy and Alzheimer disease — one peptide, two pathways. Nature Reviews Neurology 2020, 16 (1), 30–42.

5. Fotuhi, M.; Hachinski, V.; Whitehouse, P. J. Changing perspectives regarding late-life dementia. Nature Reviews Neurology 2009, 5 (12), 649–658.

6. Kalaria, R. N.; Ihara, M. Vascular and neurodegenerative pathways—will they meet? Nature Reviews Neurology 2013, 9 (9), 487–488.

7. Revesz, T.; Holton, J. L.; Lashley, T.; Plant, G.; Rostagno, A.; Ghiso, J.; Frangione, B. Sporadic and Familial Cerebral Amyloid Angiopathies. Brain Pathology 2002, 12 (3), 343–357.

8. Biffi, A.; Greenberg, S. M. Cerebral amyloid angiopathy: a systematic review. Journal of Clinical Neurology 2011, 7 (1), 1–9.

9. Plant, G. T.; RÉVÉSz, T.; Barnard, R. O.; Harding, A. E.; Gautier-Smith, P. C. Familial Cerebral Amyloid Angiopathy with nonneuritic amyloid plaque formation. Brain 1990, 113 (3), 721–747.

10. Maat-Schieman, M.; Roos, R.; van Duinen, S. Hereditary cerebral hemorrhage with amyloidosis-Dutch type. Neuropathology 2005, 25 (4), 288–97.

11. Lu, J.-X.; Qiang, W.; Yau, W.-M.; Schwieters, Charles D.; Meredith, Stephen C.; Tycko, R. Molecular Structure of β-Amyloid Fibrils in Alzheimer’s Disease Brain Tissue. Cell 2013, 154 (6), 1257–1268.

12. Tycko, R. Solid-state NMR studies of amyloid fibril structure. Annu Rev Phys Chem 2011, 62, 279–299.

13. Tycko, R. Amyloid polymorphism: structural basis and neurobiological relevance. Neuron 2015, 86 (3), 632–645.

14. Fu, Z.; Van Nostrand, W. E.; Smith, S. O. Anti-Parallel β-Hairpin Structure in Soluble Aβ Oligomers of Aβ40-Dutch and Aβ40-Iowa. International Journal of Molecular Sciences. DOI: 10.3390/ijms22031225.

15. Grant, M. A.; Lazo, N. D.; Lomakin, A.; Condron, M. M.; Arai, H.; Yamin, G.; Rigby, A. C.; Teplow, D. B. Familial Alzheimer’s disease mutations alter the stability of the amyloid β-protein monomer folding nucleus. Proceedings of the National Academy of Sciences 2007, 104 (42), 16522–16527.

16. Qiang, W.; Yau, W.-M.; Tycko, R. Structural Evolution of Iowa Mutant β-Amyloid Fibrils from Polymorphic to Homogeneous States under Repeated Seeded Growth. Journal of the American Chemical Society 2011, 133 (11), 4018–4029.

17. Qiang, W.; Yau, W.-M.; Luo, Y.; Mattson, M. P.; Tycko, R. Antiparallel β-sheet architecture in Iowa-mutant β-amyloid fibrils. Proceedings of the National Academy of Sciences 2012, 109 (12), 4443–4448.

18. Rajpoot, J.; Crooks, E. J.; Irizarry, B. A.; Amundson, A.; Van Nostrand, W. E.; Smith, S. O. Insights into Cerebral Amyloid Angiopathy Type 1 and Type 2 from Comparisons of the Fibrillar Assembly and Stability of the Aβ40-Iowa and Aβ40-Dutch Peptides. Biochemistry 2022, 61 (12), 1181–1198.

19. Selkoe, D. J.; Hardy, J. The amyloid hypothesis of Alzheimer’s disease at 25 years. EMBO Mol Med 2016, 8 (6), 595–608.

20. Chen, G.-f.; Xu, T.-h.; Yan, Y.; Zhou, Y.-r.; Jiang, Y.; Melcher, K.; Xu, H. E. Amyloid beta: structure, biology and structure-based therapeutic development. Acta Pharmacologica Sinica 2017, 38 (9), 1205–1235.

21. Dazzi, A.; Prater, C. B. AFM-IR: Technology and Applications in Nanoscale Infrared Spectroscopy and Chemical Imaging. Chemical Reviews 2017, 117 (7), 5146–5173.

22. Schwartz, J. J.; Jakob, D. S.; Centrone, A. A guide to nanoscale IR spectroscopy: resonance enhanced transduction in contact and tapping mode AFM-IR. Chemical Society Reviews 2022, 51 (13), 5248–5267.

23. Banerjee, S.; Ghosh, A. Structurally Distinct Polymorphs of Tau Aggregates Revealed by Nanoscale Infrared Spectroscopy. The Journal of Physical Chemistry Letters 2021, 12 (45), 11035–11041.

24. Banerjee, S.; Holcombe, B.; Ringold, S.; Foes, A.; Naik, T.; Baghel, D.; Ghosh, A. Nanoscale Infrared Spectroscopy Identifies Structural Heterogeneity in Individual Amyloid Fibrils and Prefibrillar Aggregates. The Journal of Physical Chemistry B 2022, 126 (31), 5832–5841.

25. Barth, A. Infrared spectroscopy of proteins. Biochimica et Biophysica Acta (BBA) – Bioenergetics 2007, 1767 (9), 1073–1101.

26. Barth, A.; Zscherp, C. What vibrations tell about proteins. Quarterly Reviews of Biophysics 2002, 35 (4), 369–430.

27. Silva, R. A. G. D.; Barber-Armstrong, W.; Decatur, S. M. The Organization and Assembly of a β-Sheet Formed by a Prion Peptide in Solution: An Isotope-Edited FTIR Study. Journal of the American Chemical Society 2003, 125 (45), 13674–13675.

28. Moran, S. D.; Zanni, M. T. How to Get Insight into Amyloid Structure and Formation from Infrared Spectroscopy. The Journal of Physical Chemistry Letters 2014, 5 (11), 1984–1993.

29. Reppert, M.; Tokmakoff, A. Electrostatic frequency shifts in amide I vibrational spectra: direct parameterization against experiment. J Chem Phys 2013, 138 (13), 134116.

30. Oh, K. I.; Fiorin, G.; Gai, F. How Sensitive is the Amide I Vibration of the Polypeptide Backbone to Electric Fields? Chemphyschem 2015, 16 (17), 3595–8.

31. Hinrichs, K.; Shaykhutdinov, T. Polarization-Dependent Atomic Force Microscopy–Infrared Spectroscopy (AFM-IR): Infrared Nanopolarimetric Analysis of Structure and Anisotropy of Thin Films and Surfaces. Applied Spectroscopy 2018, 72 (6), 817–832.

32. Antzutkin, O. N.; Balbach, J. J.; Leapman, R. D.; Rizzo, N. W.; Reed, J.; Tycko, R. Multiple quantum solid-state NMR indicates a parallel, not antiparallel, organization of beta-sheets in Alzheimer’s beta-amyloid fibrils. Proc Natl Acad Sci U S A 2000, 97 (24), 13045–13050.

33. Tycko, R. Molecular structure of amyloid fibrils: insights from solid-state NMR. Quarterly Reviews of Biophysics 2006, 39 (1), 1–55.

34. Banerjee, S.; Baghel, D.; Hasan Ul Iqbal, M.; Ghosh, A. Nanoscale Infrared Spectroscopy Identifies Parallel to Antiparallel β-Sheet Transformation of Aβ Fibrils. The Journal of Physical Chemistry Letters 2022, 13 (45), 10522–10526.

35. Sarroukh, R.; Cerf, E.; Derclaye, S.; Dufrêne, Y. F.; Goormaghtigh, E.; Ruysschaert, J. M.; Raussens, V. Transformation of amyloid β(1-40) oligomers into fibrils is characterized by a major change in secondary structure. Cell Mol Life Sci 2011, 68 (8), 1429–38.

36. Cheon, M.; Chang, I.; Mohanty, S.; Luheshi, L. M.; Dobson, C. M.; Vendruscolo, M.; Favrin, G. Structural Reorganisation and Potential Toxicity of Oligomeric Species Formed during the Assembly of Amyloid Fibrils. PLOS Computational Biology 2007, 3 (9), e173.

37. Ahmed, M.; Davis, J.; Aucoin, D.; Sato, T.; Ahuja, S.; Aimoto, S.; Elliott, J. I.; Van Nostrand, W. E.; Smith, S. O. Structural conversion of neurotoxic amyloid-beta(1-42) oligomers to fibrils. Nat Struct Mol Biol 2010, 17 (5), 561–7.

