## Supplementary Information for "Intermediate antiparallel fibrils in Aβ40 Dutch mutant aggregation: nanoscale insights from AFM-IR"

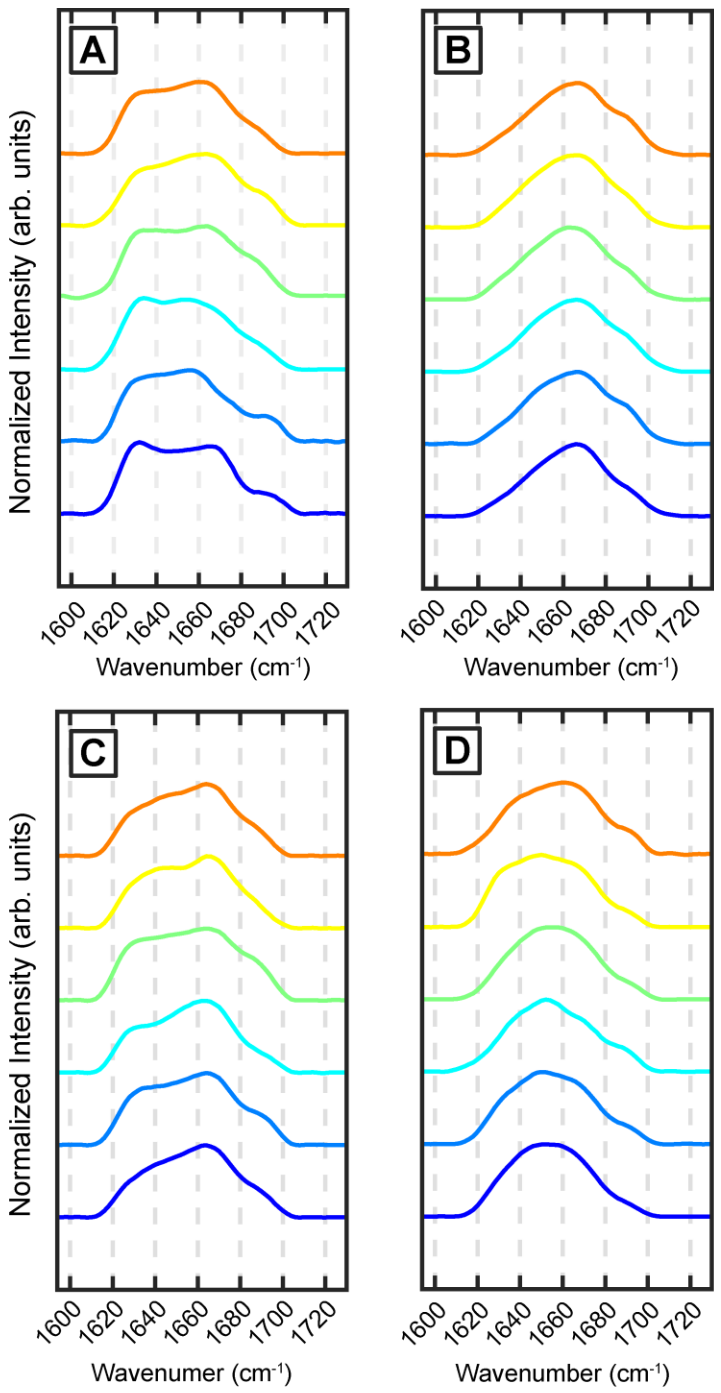


**Fig S1.** Representative IR spectra obtained from Aβ40 (E22Q) Dutch mutant oligomers. Two types of amide I bands have been observed. (A) Type 1 oligomer with more parallel β sheet content and (B) type 2 oligomers having less parallel β sheet have been identified after 2 h of aggregation. (C-D) Similar two types of oligomers are found after 8 h of aggregation.

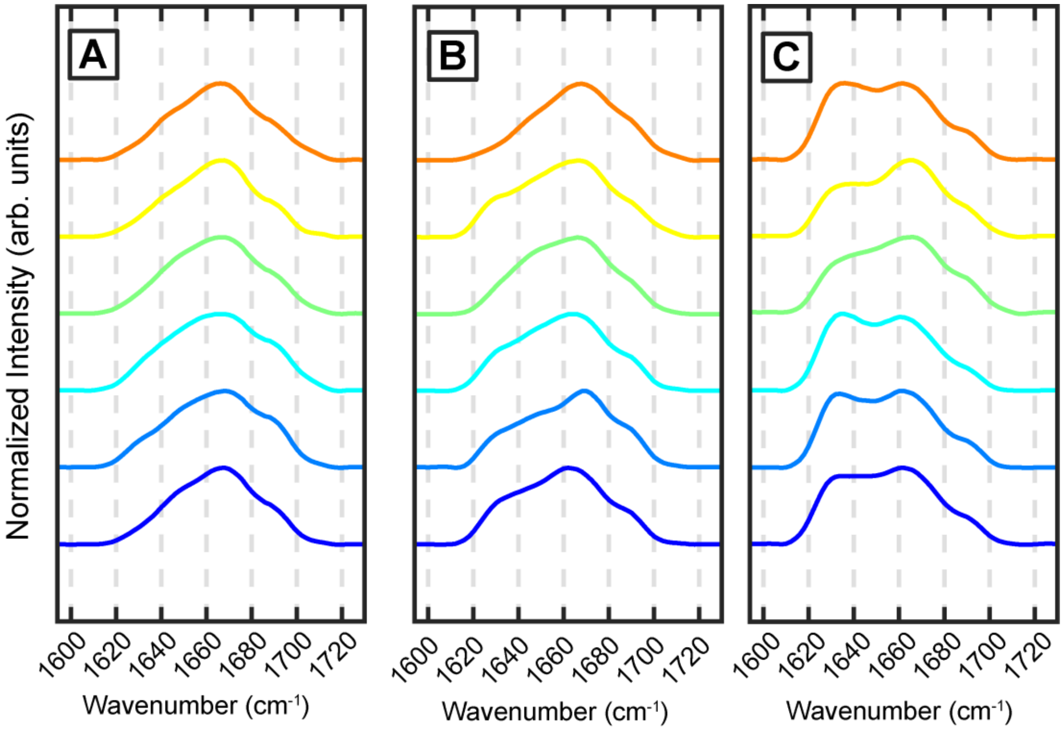


**Fig S2.** Representative IR spectra obtained from Aβ40 (E22Q) Dutch mutant fibrils generated after (A) 36 h, (B) 72 h and (C) 8 days.

**Spectral Fitting of spectra:**

The mean spectra shown in Figures 1-2 in the main manuscript were fit to a sum of four constituent Gaussian peaks:

$$S = \sum_{n=1}^{4} A_n e^{-\left(\frac{\omega - \omega_{0,n}}{\sigma}\right)^2}$$

where $A_n$ is the amplitude of the n-th peak, and $\omega_{0,n}$ the corresponding center frequency. The spectra were normalized to the maximum intensity of the amide-I band prior to fitting. The peak positions identified in the second derivative spectra, shown in Figures 1-2 in the manuscript, were used as a starting point for the fitting procedure. The center frequency was allowed to vary between ±5cm$^{-1}$ of the initial value.

**Supplementary Table 1. Fitting parameters**

| | Center frequency ($\omega_0$, cm$^{-1}$) | Amplitude (A) | Width (σ, cm$^{-1}$) |
|---|---|---|---|
| 2-h fibril type-1 | 1628.0 | 0.50 | 9.46 |
| | 1640.0 | 0.54 | 13.74 |
| | 1663.0 | 0.96 | 20.0 |
| | 1692.0 | 0.17 | 8.04 |

| | Center frequency ($\omega_0$, cm$^{-1}$) | Amplitude (A) | Width (σ, cm$^{-1}$) |
|---|---|---|---|
| 2-h fibril type-2 | 1629.0 | 0.10 | 9.24 |
| | 1645.0 | 0.43 | 15.39 |
| | 1668.0 | 0.95 | 19.44 |
| | 1691.0 | 0.20 | 9.05 |

| | Center frequency ($\omega_0$, cm$^{-1}$) | Amplitude (A) | Width (σ, cm$^{-1}$) |
|---|---|---|---|
| 8-h fibril type-1 | 1628.0 | 0.40 | 9.10 |
| | 1642.0 | 0.58 | 14.67 |
| | 1666.0 | 0.95 | 19.20 |
| | 1691.0 | 0.21 | 8.05 |

| | Center frequency ($\omega_0$, cm$^{-1}$) | Amplitude (A) | Width (σ, cm$^{-1}$) |
|---|---|---|---|
| 8-h fibril type-2 | 1630.0 | 0.24 | 9.64 |
| | 1646.0 | 0.76 | 15.01 |
| | 1667.0 | 0.83 | 16.13 |
| | 1691.0 | 0.16 | 8.30 |

| | Center frequency ($\omega_0$, $cm^{-1}$) | Amplitude (A) | Width (σ, $cm^{-1}$) |
|---|---|---|---|
| 36-h | 1630.0 | 0.18 | 8.90 |
| | 1646.0 | 0.55 | 14.97 |
| | 1668.0 | 0.92 | 16.8 |
| | 1692.0 | 0.31 | 8.66 |

| | Center frequency ($\omega_0$, $cm^{-1}$) | Amplitude (A) | Width (σ, $cm^{-1}$) |
|---|---|---|---|
| 72-h | 1628.0 | 0.20 | 9.13 |
| | 1643.0 | 0.50 | 14.83 |
| | 1667.0 | 0.96 | 18.38 |
| | 1691.0 | 0.23 | 8.50 |

| | Center frequency ($\omega_0$, $cm^{-1}$) | Amplitude (a) | Width (σ, $cm^{-1}$) |
|---|---|---|---|
| 8 days | 1628.0 | 0.30 | 8.52 |
| | 1638.0 | 0.47 | 12.29 |
| | 1665.0 | 0.99 | 20 |
| | 1691.0 | 0.21 | 7.71 |
